# A near-complete telomere-to-telomere genome assembly for *Batrachochytrium dendrobatidis* GPL JEL423 reveals a CBM18 gene family expansion and a M36 metalloprotease gene family contraction

**DOI:** 10.1101/2024.10.22.619730

**Authors:** Nicolas Helmstetter, Keith Harrison, Jack Greggory, Jamie Harrison, Elizabeth Ballou, Rhys A. Farrer

## Abstract

*Batrachochytrium dendrobatidis* (*Bd*) is responsible for mass extinctions and extirpations of amphibians, mainly driven by the Global Panzootic Lineage (*Bd*GPL). *Bd*GPL isolate JEL423 is a commonly used reference strain in studies exploring the evolution, epidemiology and pathogenicity of chytrid pathogens. These studies have been hampered by the fragmented, erroneous and incomplete *B. dendrobatidis* JEL423 genome assembly, which includes long stretches of ambiguous positions, and poorly resolved telomeric regions. Here we present and describe a substantially improved, near telomere-to-telomere genome assembly and gene annotation for *B. dendrobatidis* JEL423. Our new assembly is 24.5 Mb in length, ∼800 kb longer than the previously published assembly for this organism, comprising 18 nuclear scaffolds and 2 mitochondrial scaffolds and including an extra 839 kb of repetitive sequence. We discovered that the patterns of aneuploidy in *B. dendrobatidis* JEL423 have remained stable over approximately 5 years. We found that our updated assembly encodes fewer than half the number of M36 metalloprotease genes predicted in the previous assembly. In contrast, members of the crinkling and necrosis gene family were found in similar numbers to the previous assembly. We also identified a more extensive carbohydrate binding module 18 gene family than previously observed. We anticipate our findings, and the updated genome assembly will be a useful tool for further investigation of the genome evolution of the pathogenic chytrids.

**Author Summary:** *B. dendrobatidis* is a fungus that has been implicated in the extinctions and declines of dozens of amphibians globally. Here we describe a new genome assembly for the commonly used *B. dendrobatidis* isolate JEL423, which is a substantial improvement from the previously used assembly. Compared with the previous assembly, we reveal that some gene families (such as carbohydrate binding module 18 genes) are more expanded than previously recognised, while others (such as the M36 metalloproteases) are more contracted than previously recognised. Our new genome assembly will be useful for future work exploring the pathogenicity, epidemiology and evolution of chytrid fungi.

## Introduction

Over 40% of amphibians are threatened with extinction (The IUCN Red List of Threatened Species 2022), attributed to several factors including pollution, habitat loss and infectious diseases including the globally distributed chytrid fungus *Batrachochytrium dendrobatidis* (Longcore et al. 1999; Farrer 2019). *B. dendrobatidis* is thought to have caused the extinction of over 90 species to date, with a further 500 in decline (Scheele et al. 2019). *B. dendrobatidis* causes chytridiomycosis, which degrades the skin of amphibians and the keratinized mouthparts of tadpoles. Several putative virulence factors and genes involved in pathogenicity have been identified from genomic studies of *B. dendrobatidis*, including the expanded family of M36 metalloproteases thought to degrade host skin and extracellular matrix (Joneson et al. 2011; Farrer et al. 2017a; Wacker et al. 2023), the enigmatic expanded family of genes with sequence similarity to crinkling and necrosis (CRN) genes (Amaro et al. 2017; Farrer et al. 2017a), as well as carbohydrate binding module 18 gene family (CBM18) thought to bind and thereby limit exposure of fungal chitin (Liu and Stajich 2015; Farrer et al. 2017a). Several gene family expansions can be observed in the previous *B. dendrobatidis* genome assembly, including that of the M36 metalloproteases, these have been linked to transposon proliferation and evolutionary selection (Wacker et al. 2023). Furthermore, loss of heterozygosity (LOH) and whole chromosome aneuploidy have been detected in *B. dendrobatidis* genomes, possibly linked to its pathogenic lifestyle (Farrer et al. 2013). However, the specific genes and mechanisms underlying *B. dendrobatidis* pathogenicity remain poorly understood, owing to a lack of experimental validation of these pathogenicity factors and dated genomic resources.

*B. dendrobatidis* has genetically diverse populations, comprising 5 known lineages to date: The Global Panzootic Lineage (*Bd*GPL), *Bd*ASIA-1, *Bd*ASIA-2, *Bd*ASIA-3, and *Bd*CAPE (Farrer et al. 2011a; O’Hanlon et al. 2018; Wacker et al. 2024). Of the five known lineages, *Bd*GPL is characterised as hypervirulent and globally distributed (Fisher and Garner 2020) and is thought to be the main driver of the chytridiomycosis panzootic (Farrer et al. 2011b). *Bd*GPL isolate JEL423 was originally isolated by Dr. Joyce Longcore at the University of Maine from a lemur leaf frog (*Agalychnis lemur*) in Panama. *B. dendrobatidis* JEL423 was sequenced and assembled in 2006 into 69 scaffolds with a total length of 23.7 Mb, using a hybrid approach including Sanger and Illumina data (no manuscript accompanied this data, although the revised annotation and polished assembly is described here (Farrer et al. 2017b)).

In 2017, the assembly was updated and improved (version 2 of the assembly) using Illumina DNAseq based Pilon polishing and gene annotation refinement using RNA-seq (Farrer et al. 2017b). In 2018 the structure of the mitochondrial genome was resolved using long read sequencing, into three linear segments (O’Hanlon et al. 2018). Here we describe an updated, near telomere-to-telomere genome assembly and gene annotation for *B. dendrobatidis* JEL423 (version 3) that will be a crucial resource for understanding *B. dendrobatidis*’ evolution to a pathogenic lifestyle.

## Results

We re-assembled *B. dendrobatidis* JEL423 using long Oxford Nanopore Technologies (ONT) and Hi-C sequencing, which substantially improved assembly metrics compared with the previous V2 assembly (**Fig. 1**; **Table 1**). Specifically, the V2 23.7 Mb assembly is highly fragmented (69 scaffolds, 348 contigs) and includes many long stretches of ambiguous bases (318,766 Ns in total), while our new assembly is ∼800 kb longer (24.5 Mb), comprising just 18 nuclear scaffolds, 2 mitochondrial scaffolds, 22 contigs and only 4 gaps. The final Hi-C contact map (**Fig. 1B**) validates the contiguity of the assembly, showing the strong self-interactions of scaffolds 1 to 16 and the clear junctions between each of them, typical of chromosomes. Those chromosomes seem to be organised in a Rabl-like architecture with prominent contacts between telomeres (Hoencamp et al. 2021).

**Figure 1.**
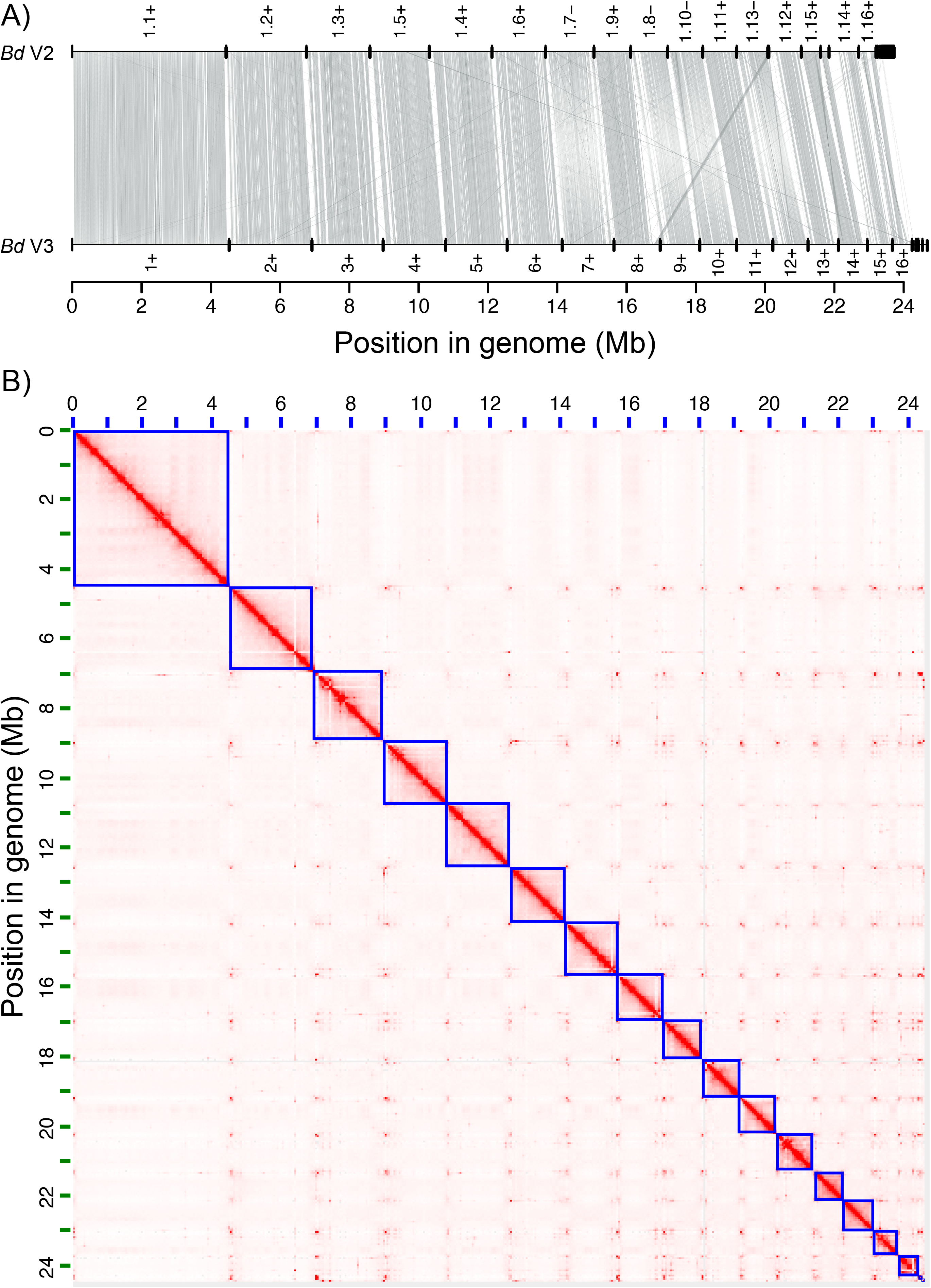
(**A**) Synteny plot of previous assembly and annotation for *B. dendrobatidis* JEL423 (V2) compared with the new version (V3). Scaffold numbers are shown, along with the orientation. (**B**) A chromatin capture contact map of *B. dendrobatidis*, where blue squares indicate putative chromosomes, and bolder red indicates increased contact frequency. The Rabl-like phenomenon found in other species, where telomeres co-situate in the nucleus, is visible.

**Table 1.** Genome assembly metrics of the previous *B. dendrobatidis* assembly (V2) compared with our updated genome assembly (V3).

| Genome assembly | JEL423 (V2) | JEL423 (V3) |
| --- | --- | --- |
| Total Length (Mb) | 23.7 | 24.5 |
| Contig count | 348 | 22 |
| Contig N<sub>50</sub> (Mb) | 0.22 | 1.49 |
| Contig N<sub>90</sub> (Kb) | 42 | 752 |
| Contig L<sub>50</sub> | 32 | 6 |
| Contig L<sub>90</sub> | 129 | 15 |
| Scaffold count | 69 | 18 |
| Scaffold N<sub>50</sub> (Mb) | 1.70 | 1.77 |
| Scaffold N<sub>90</sub> (Kb) | 857 | 877 |
| Scaffold L<sub>50</sub> | 5 | 5 |
| Scaffold L<sub>90</sub> | 14 | 13 |
| Number of gaps | 279 | 4 |
| Ns per 100kb | 13.44 | 0.02 |
| Largest scaffold (Mb) | 4.4 | 4.5 |
| Telomeres total | 0 | 18 |
| Telomeres forward (5') | 0 | 8 |
| Telomeres reverse (3') | 0 | 10 |
| Telomere to telomere scaffolds | 0 | 5 |
| GC content (%) | 39.29 | 39.28 |
| Protein coding genes | 8630 | 8628 |
| BUSCO complete (%) | 91.4 | 92.1 |
| Short read alignment rate (%) | 94.36 | 99.03 |
| Long reads alignment rate (%) | 93.27 | 96.17 |

To verify the accuracy of the improved genome assembly we calculated mapping rates by aligning Illumina data (O’Hanlon et al. 2018) and long reads (this study) to the genome assembly. As a result, 99.02% of the Illumina reads aligned (up from 94.36% for *B. dendrobatidis* V2) and 96.17% of the long reads aligned (up from 93.27% for *B. dendrobatidis* V2). Furthermore, the V2 assembly had no contigs ending in telomeric repeats, while our updated assembly is near telomere-to-telomere, with 13/18 (72%) scaffolds terminating with 1 or 2 telomeric repeat (TTAGGG), while 5/18 (28%) have telomeres at both ends (**Table S1**). Finally, our new, more contiguous mitochondrial genome assembly is ∼ 260 kb: a reduction from the previous assembly of nearly 300 kb.

We re-annotated our assembly, which identified 8,628 predicted protein coding genes (excluding splice variants), which was very similar to previously predicted (*n* = 8,630). Benchmarking Universal Single-Copy Orthologs (BUSCO) scores from the new assembly were slightly improved compared with the V2 assembly (92.1% up from 91.4%), suggesting a higher overall accuracy in assembly and gene prediction. Our improved genome assembly demonstrated many genomic rearrangements compared to the previous assembly, in addition to the increased overall length and contiguity (**Fig. 1A**). Comparing the new *B. dendrobatidis* assembly to genome assemblies from its closest-known chytrid relatives supported the relationships and the low-level of conserved synteny previously identified between those species, as well as highlighting the fragmented and low-quality assemblies of those other species being compared to (**Fig. 2**).

**Figure 2.**
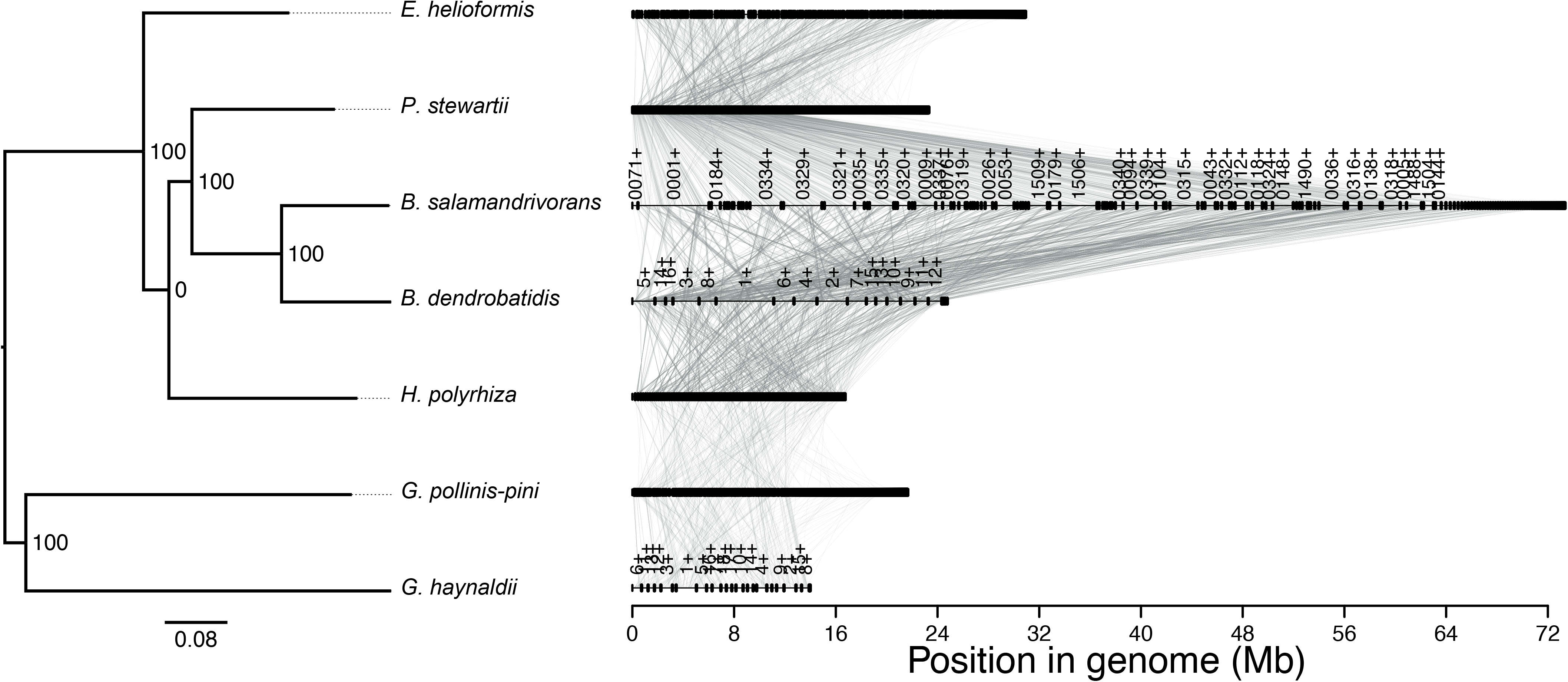
A synteny plot between the new *B. dendrobatidis* genome assembly and its closest relatives. Synteny is poorly conserved between the species represented. Additionally, the genome assemblies of all species apart from our new *B. dendrobatidis* JEL423 assembly reported here remain fragmented or highly fragmented.

Our new assembly facilitated the prediction of a further 839 kb of repetitive sequence compared with the previous assembly (23.89% instead of 21.42%; **Table 2**). Notably, increases in the numbers of unclassified and total interspersed repeats, long terminal repeat (LTR) elements, DNA transposons and retroelements, as well as small RNAs were identified in the new assembly. Mutator, Cacta and Maverick-type DNA transposons are the most common in the *B. dendrobatidis* genome assembly followed closely by retrotransposons (**Table S2**). Conversely, there were small decreases in the numbers of short interspersed nuclear elements, long interspersed nuclear elements, and low complexity repeats.

**Table 2.** Repeat content predicted from the previous *B. dendrobatidis* assembly (V2) compared with our updated genome assembly (V3) based on RepeatMasker and RepeatModeller.

|  | JEL423 (V2) |  |  | JEL423 (V3) |  |  |  |
| --- | --- | --- | --- | --- | --- | --- | --- |
|  | number | length (bp) | genome occupied (%) | number | length (bp) | genome occupied (%) | Difference (nt) |
| **Total (bases masked)** |  | 5,013,710 | 21.42 |  | 5,852,638 | 23.89 | 838,928 |
| **Retroelements** | 265 | 267,526 | 1.14 | 513 | 297,044 | 1.21 | 29,518 |
| **SINEs** | 45 | 30,456 | 0.13 | 28 | 17,579 | 0.07 | -12,877 |
| **LINEs** | 108 | 152,775 | 0.65 | 62 | 129,726 | 0.53 | -23,049 |
| **LINEs (R1/LOA/Jockey)** | 19 | 12,358 | 0.05 | 0 | 0 | 0 | -12,358 |
| **LTR elements** | 112 | 84,295 | 0.36 | 423 | 149,739 | 0.61 | 65,444 |
| **LTR elements (BEL/Pao)** | 0 | 0 | 0 | 322 | 52,204 | 0.21 | 52,204 |
| **LTR elements (Ty1/Copia)** | 63 | 63,588 | 0.27 | 75 | 86,851 | 0.35 | 23,263 |
| **LTR elements (Gypsy/DIRS1)** | 33 | 10,112 | 0.04 | 26 | 10,684 | 0.04 | 572 |
| **LTR elements (Gypsy/DIRS1 - Retroviral)** | 16 | 10595 | 0.05 | 0 | 0 | 0 | -10,595 |
| **DNA transposons** | 565 | 775,071 | 3.31 | 719 | 804,803 | 3.28 | 29,732 |
| **Unclassified** | 9,174 | 3,910,035 | 16.71 | 9,409 | 4,448,145 | 18.15 | 538,110 |
| **Total interspersed repeats** |  | 4,952,632 | 21.16 |  | 5,549,992 | 22.65 | 597,360 |
| Small RNA | 0 | 0 | 0 | 50 | 234,327 | 0.96 | 234,327 |
| Simple repeats | 1,191 | 54,799 | 0.23 | 1,310 | 63,418 | 0.26 | 8,619 |
| Low complexity | 115 | 6,279 | 0.03 | 99 | 4,901 | 0.02 | -1,378 |

The ploidy of *B. dendrobatidis* JEL423 was assessed using previously generated Illumina reads (O’Hanlon et al. 2018) for the isolate aligned to the new assembly, revealing widespread aneuploidy and high base ploidy level for this isolate as previously described (Farrer et al. 2013) (**Fig. 3**). For example, normalised depth showed differences between scaffolds, with scaffolds 2, 3, 8, 10, 11, 13, 15 and 16 all at different levels to scaffold 1, and about 4 separate depth levels identified (**Fig. 3A**). There were also dips in normalised depth at various points within scaffolds 1, 2, 3, 4, 6 and 10 suggestive of centromeric regions.

**Figure 3.**
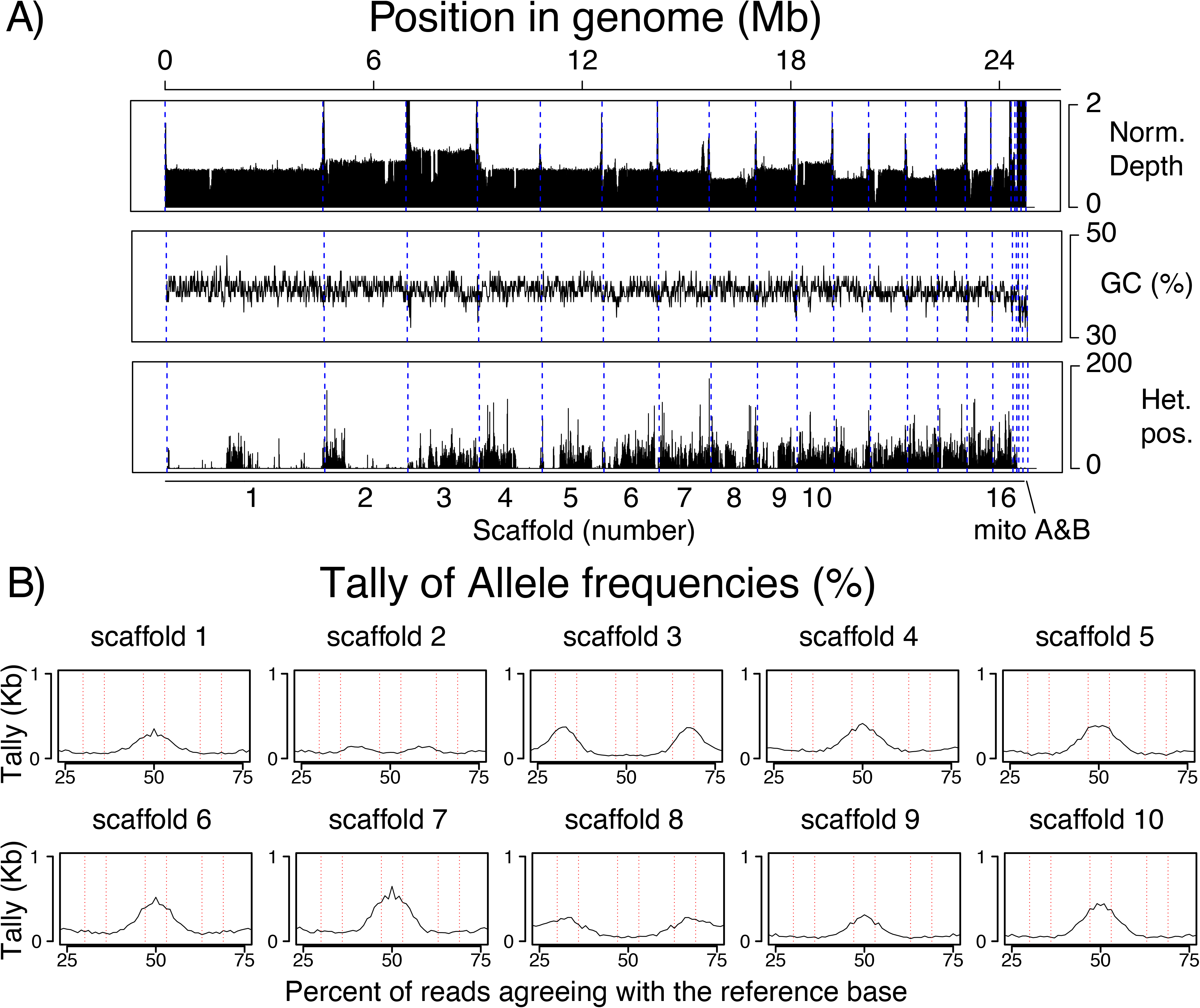
Ploidy of *B. dendrobatidis* JEL423 was assessed, based on (**A**) normalised depth of coverage across 10 kb non-overlapping windows, GC content and tallying of heterozygous positions across the genomes, and (**B**) allele frequencies, where the percent of reads agreeing with the reference base was counted and tallied across every base in each chromosome, and the percents between 25% and 75% plotted. These analyses support previous findings that *B. dendrobatidis* JEL423 has widespread aneuploidy, including diploidy, triploidy, and perhaps tetraploidy+. Furthermore, loss of heterozygosity is seen across the genome, particularly in scaffolds 1, 2 and 4.

While GC content is relatively stable across the genome, LOH is stark, with several genomic regions showing an enrichment or dearth of heterozygosity. For example, the middle of scaffold 1, the start of scaffold 2 and half of scaffold 4 have high levels of heterozygosity, while the majority of scaffold 1, and ends of scaffolds 2 and 4 are particularly lacking in heterozygosity (**Fig. 3A**).

We counted the percent of reads agreeing with the reference base across every scaffold to help predict ploidy (**Fig. 3B**). Most positions had ∼ 100% of reads agreeing (reference/homozygous positions), and a smaller number had ∼ 0% of reads agreeing (SNP/homozygous or assembly errors) as expected, neither of which are shown in **Fig. 3B**. Many genomic positions indicative of heterozygosity were also identified (**Fig. 3B**). For example, scaffold 1 has a bi-allelic peak suggestive of an even ploidy (such as diploid or tetraploid), scaffold 2 has slightly higher depth and ambiguous or little evidence of heterozygosity (owing in part to the LOH for most of that scaffold), scaffold 3 has higher depth than scaffolds 1 and 2 and has peaks at 33% and 67% indicative of an odd ploidy of triploid or higher, while scaffold 4 returns to scaffold 1 depth of coverage and has a bi-allelic peak suggestive of an even ploidy. Scaffold 8 (along with scaffolds 11 and 13) has the lowest normalised depth level (**Fig. 3A**) and Scaffold 8 shows trimodal heterozygosity peaks suggesting triploid, which together with the above results, suggest *B. dendrobatidis* JEL423 is base tetraploid, with some triploid and pentaploid scaffolds. More surprising, is that the pattern of normalised depth and allele frequencies is very consistent with the previous assemblies and the Depth of coverage and allele frequencies identified from our 2013 paper (Farrer et al. 2013) and 2018 paper (O’Hanlon et al. 2018), with sequencing (and culturing) occurring approximately 5 years apart, indicating that this aneuploidy is not transient as previously speculated (Farrer et al. 2013), but may be stable, at least across several years in passage and/or cryogenically frozen stocks.

We found a reduction in the number of encoded M36 metalloprotease, which are a key predicted pathogenicity factor, in the new assembly (*n* = 14) compared with the old assembly (*n* = 37) (**Fig. 4**). Indeed, we found fewer predicted proteases overall (*n* = 722) compared with the previous assembly (*n* = 804) (**Fig. S1**). To ensure the reduction in number of M36 metalloprotease genes was not a result of erroneous gene calling, we used tblastn of all our new M36 genes to the genome assembly, identifying 8 further genes with sequence similarity (**Fig. S2**). However, only one of these newly identified genes had predicted protease function (A1 protease g2465). The majority of the newly identified M36 genes belonged to an orthogroup with M36 in the V2 assembly (*n* = 10/14). The newly identified M36 genes fall within the *B. dendrobatidis* and *B. salamandrivorans* metalloprotease clade as previously identified (**Figs. 4B** and **4C**). However, the reduced number in *B. dendrobatidis* suggests the gene expansion of M36s in *B. salamandrivorans* is even more substantial than previously recognised.

**Figure 4.**
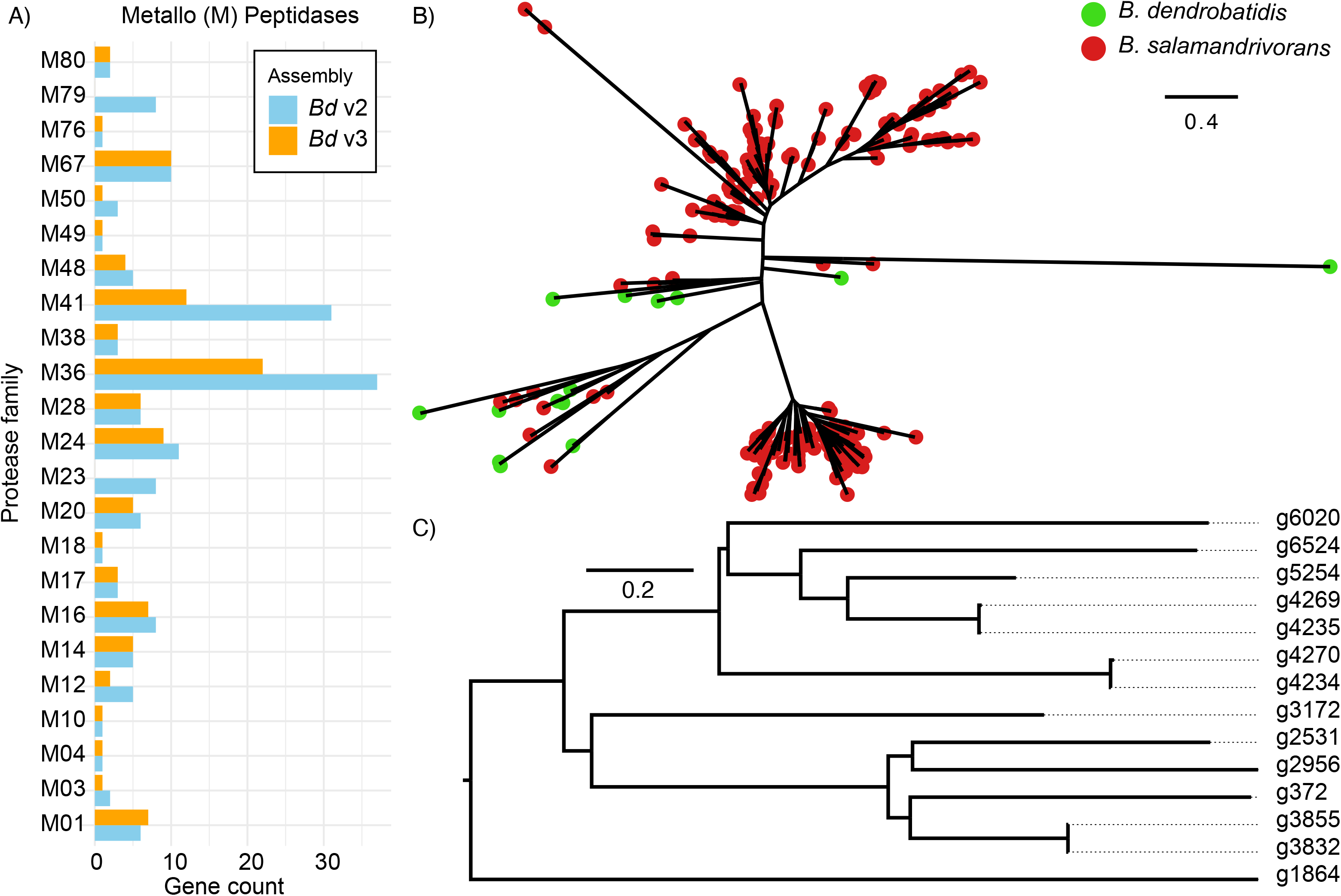
Predicted M36 metalloproteases in *B. dendrobatidis* JEL423. (**A**) metalloproteases had a modest difference in total count between the old (V2) and new (V3) assemblies, with substantial decreases in the number of M36 and M41 proteases. (**B**) A gene tree (FastTree) between *B. dendrobatidis* JEL423 (V3) and *B. salamandrivorans* M36’s supports previous findings of multiple sub gene families and ancestral gene expansions, particularly in *B. salamandrivorans* since its split with *B. dendrobatidis*. (**C**) A gene tree (FastTree) of predicted *B. dendrobatidis* M36’s. Branch lengths indicate the mean number of nucleotide substitutions per site.

Unlike the M36 metalloproteases, a large expansion of CRN genes are still predicted as a unique feature of the *B. dendrobatidis* genome. Using BLASTp for all proteins against the CRNs predicted in *Phytophthora infestans* T30-4 (where CRNs are better described) as previously performed (Farrer et al. 2017b), revealed only 13 top high scoring pairs (HSPs) after excluding splice variants (**Fig. S3**). However, after running tblastn against the genome, 133 separate putative CRNs were identified, which is similar to the number previously identified in the V2 assembly (*n* = 162; 82%) (**Fig. 5**).

**Figure 5.**
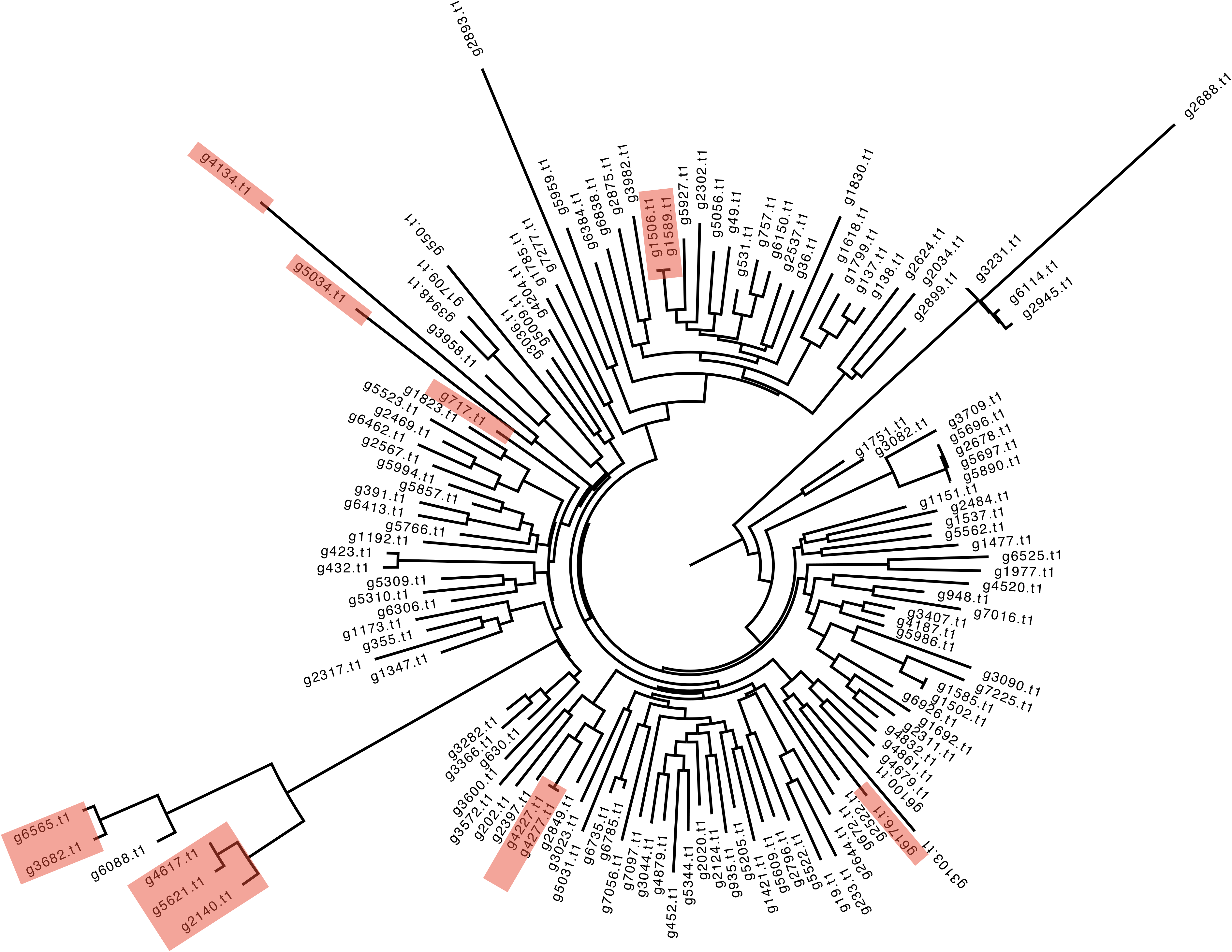
Predicted crinkling and necrosis (CRN) genes in *B. dendrobatidis* JEL423 based on a BLASTp search to the *P. infestans* T30-4 as previously done (Farrer et al. 2017b), revealed only 13 top HSPs after excluding splice variants (highlighted in red), along with 120 further genes identified from tblastn. The gene tree was constructed using FastTree. Branch lengths indicate the mean number of nucleotide substitutions per site.

The CBM18 gene family has nearly double the predicted genes in the updated assembly compared with the V2 assembly (n = 39 compared with 22) (**Fig. 6**). Based on BLASTp and HMMER searches, the number of Tyrosine-like increased from 5 to 12, the number of Deacetylase-like increased from 10 to 19, and the number of lectin-like remained at 5. While the Tyrosine-like and Deacetylase-like were easy to predict based on PFAM domains, the lectin-like were less clear, as these were based on previous predictions, and the BLASTp searches overlapped with Deacetylase-like genes. Notably, gene g231 has the lowest BLASTp e-value (e-value = 0, compared with the next lowest at 1.09e-51 for g6713 which is clustered with, and contains a Tyrosine-like PFAM domain).

**Figure 6.**
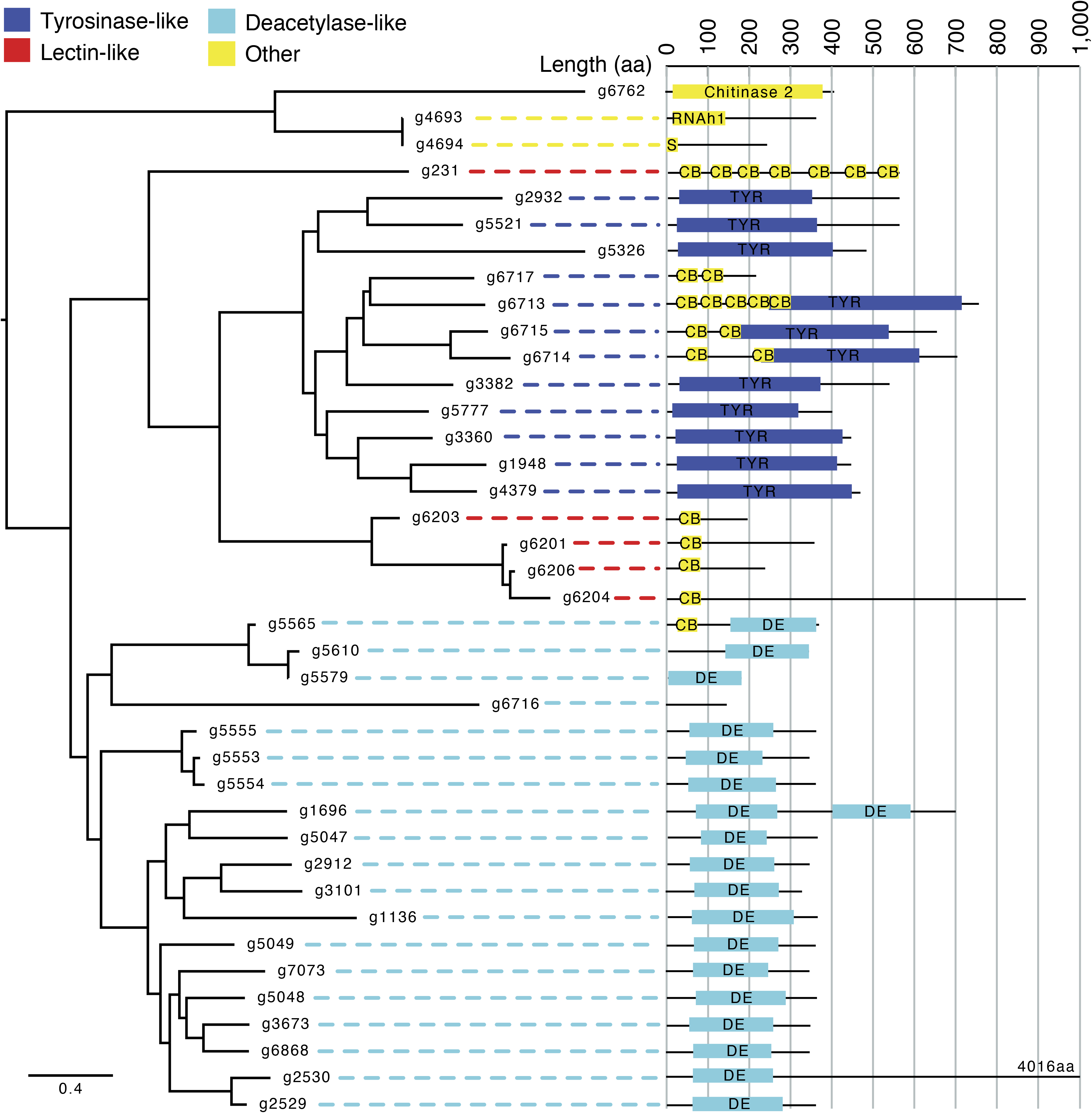
Predicted CBM18 genes in *B. dendrobatidis* JEL423. A phylogenetic tree of all CBM18 proteins based on BLAST to previously identified CBM18s and HMMER to PF00187, aligned using MUSCLE and a tree constructed with FastTree (branch lengths indicate the mean number of nucleotide substitutions per site). To the right of gene names is a diagram of the domain structure (RNAh1 = RNase H type-1, S = Secretion Signal, CB = Carbohydrate Binding, TYR = Tyrosinase copper-binding domain, DE= Glycoside hydrolase/deacetylase). Lectin-like are based on sequence similarity (BLASTp) to previously identified proteins (Liu and Stajich 2015).

## Discussion

*B. dendrobatidis* is responsible for mass extinctions and extirpations of amphibians. *B. dendrobatidis* isolate JEL423 was initially isolated from a lemur leaf frog in Panama during a notable extinction event across Central America, and the genome was assembled in 2006 (Farrer et al. 2017b). Since then, the reference genome of *B. dendrobatidis* JEL423 has been widely used to help understand, track and predict the species evolution, epidemiology and pathogenicity (Farrer et al. 2011b; Joneson et al. 2011; Abramyan and Stajich 2012; Liu and Stajich 2015; Farrer et al. 2017b; O’Hanlon et al. 2018; Wacker et al. 2023; Wacker et al. 2024). Five lineages of *B. dendrobatidis* have been described to date (Farrer et al. 2011b; Rosenblum et al. 2013; O’Hanlon et al. 2018), and reference genomes have been assembled for two of these lineages (*Bd*GPL and *Bd*Brazil), with recent efforts revealing pan-genomic variation in gene family counts (Yacoub and Stajich 2024). These efforts have been hampered by the fragmented, erroneous and incomplete *B. dendrobatidis* JEL423 assembly, which includes long stretches of ambiguous positions and particularly poorly resolved telomeric regions. Here, we present and describe a substantially improved, near telomere-to-telomere genome assembly and gene annotation for *B. dendrobatidis* JEL423, which will be a crucial resource for future efforts to understand *B. dendrobatidis*’ evolution to a pathogenic lifestyle.

Our new assembly is ∼800 kb longer (24.5 Mb) and comprises an extra ∼2.5% of repetitive sequence, with particular increases in unclassified and total interspersed repeats, LTR elements, DNA transposons and retroelements, which have been recently shown to associate and potentially contribute to the expansion of pathogenicity genes in the batrachochytrids (Wacker et al. 2023; Wacker et al. 2024). Mutator transposons are the most commonly identified DNA transposon in *dendrobatidis*, which are known to induce high mutation rates in other genomes and could drive genetic diversity and enhance pathogenicity (Dupeyron et al. 2019). Many Cacta elements were identified across the *dendrobatidis* genome, which are associated with chromosomal rearrangements (Badet et al. 2024), while Copia retrotransposons identified can alter gene expression (Gao et al. 2008; Muszewska et al. 2011). Finally, Maverick elements were also common in *B. dendrobatidis*, and are tied to viral function possibly contributing to genome defence and virulence factors (Kapitonov and Jurka 2006). Further work will be required to analyse and understand the distributions of repeats in the new assembly and particularly their associations with genes of interest including the M36 metalloproteases.

*B. dendrobatidis* JEL423 harbours an aneuploid genome, although it is unclear if this is found across multiple chytrid life stages, owing to the mixed cultures used for sequencing. We identified abundant heterozygosity, along with large stretches of homozygosity consistent with loss of heterozygosity events that may be the result of a hybrid origin for *Bd*GPL and/or parasex and mitotic recombination, supported further by hybrid genotypes being discovered (Farrer et al. 2011b; Greenspan et al. 2018; O’Hanlon et al. 2018; Farrer 2021). We hypothesise that the base-ploidy of *B. dendrobatidis* JEL423 is tetraploid, based on scaffolds with a decreased depth of coverage coinciding with bi-modal allele frequencies peaking at 33% and 66% agreement of reference bases – which would suggest those are triploid. Indeed, other scaffolds increase in depth and have allele frequencies consistent with pentaploidy. Surprisingly, the specific counts of chromosomes and pattern of aneuploidy appears to be stable in culture and/or cryogenically frozen stocks over several years in *B. dendrobatidis* JEL423, based on the consistency of our new results and depth of coverage across the previous assemblies and ∼ 5 year old Illumina sequencing run (Farrer et al. 2011b; Farrer et al. 2013). Further, potentially long-term evolutionary work will be required to check how long such aneuploidy can remain stable for.

Our new gene annotation revisited the counts of important gene families described previously in *B. dendrobatidis*, namely M36 metalloproteases, CRNs and CBM18s. We surprisingly identified fewer than half the number of M36 metalloprotease genes previously predicted, suggesting that the gene expansion in *B. dendrobatidis* was a result of incorrectly identified duplications in the previous assembly. The reduced number suggests that a smaller number of genes may be responsible for the destruction of skin and extracellular matrix in amphibians, and/or that other genes and gene families have a larger role than previously thought, such as the aspartyl proteases and other secreted protein families encoded (Farrer et al. 2017b). In contrast, CRNs remained abundant, albeit also slightly reduced in number, and remain enigmatic in function and evolutionary origin. CBM18s thought to bind and thereby limit exposure of fungal chitin (Liu and Stajich 2015; Farrer et al. 2017a) were also found in greater numbers than previously recognised. The placement of one predicted CBM18 (g231) in our gene tree suggest that Tyrosine-like CBM18s arose from lectin-like genes, albeit based on only preliminary and ad-hoc gene classifications.

*B. dendrobatidis* remains a substantial challenge for ecosystem health and amphibian conservation efforts (Scheele et al. 2019). Our new genome assembly will be a useful tool for further investigation the genome evolution of the pathogenic chytrids, including deciphering genomic rearrangements, gene family changes (expansions and contractions) and pan-genomics. We also anticipate that the genome assembly will help inform the design of genetic transformation approaches to explore gene function and chytrid cell biology, such as those recently developed (Kalinka et al. 2024; Webb et al. 2024). High quality genome assemblies such as our new *B. dendrobatidis* JEL423 assembly are urgently required in mycoinformatic studies, for which many species still lack a genome assembly at all, or have only draft assemblies, that can result in inaccurate descriptions of the genes and repeat-content that may be pivotal for understanding their evolution and pathogenicity.

## Methods

### DNA extraction and sequencing

*B. dendrobatidis* strain JEL423 zoosporangia and zoospores were cultured in half strength tryptone gelatin hydrolysate lactose (TGhL) (0.8% tryptone, 0.1% gelatin hydrolysate, 0.2% lactose) broth in static cell culture flasks at 18°C. High molecular weight DNA was obtained by a customized cetyltrimethylammonium bromide (CTAB) extraction procedure (Schwessinger 2019 May 17). Briefly, cell pellet was ground with a mortar and pestle in liquid nitrogen, incubated in a CTAB lysis buffer with proteinase K (460 units/mL) and RNAse A (570 units/mL), followed by potassium acetate precipitation, phenol chloroform isoamyl alcohol purification and isopropanol precipitation. Extracted DNA was checked for integrity on agarose gel and for purity and concentration with Nanodrop and Qubit. After the first extraction, 260/230 absorbance was 1.62 and the ratios between concentrations measured with qubit and nanodrop was 1/8. DNA was further purified with a QIAgen Plant Pro column resulting in 260/230 absorbance of 2.82 and concentrations ratio of 1.

An ONT library was prepared by the Exeter Sequencing Service with kit SQK- LSK109 (Oxford Nanopore Technologies) following manufacturer’s instructions. Briefly, DNA ends were FFPE repaired and end-prepped/dA-tailed using the NEBNext FFPE DNA Repair kit (M6630, NEB) and the NEBNext Ultra II End-Repair/dA-tailing Module (E7546, NEB) followed by AMPure XP bead clean-up (A63882, Beckman Coulter). Adapters were ligated using the Genomic DNA by Ligation kit (SQK-LSK109, Oxford Nanopore Technologies) and NEBNext Quick T4 DNA Ligase (E6056, NEB) followed by AMPure XP bead clean-up. The library was loaded on a Flongle flow cell (FLO-FLG001). Sequencing run yielded 1.1Gb with *N_50_* of 19.4kb. Basecalling was achieved using dorado v0.5.0 (0.5.0+0d932c0) (https://github.com/nanoporetech/dorado) with the arguments “--recursive --device=’cuda:1’ --emit-fastq dna_r9.4.1_e8_sup@v3.6 ../fast5”.

A Hi-C library was constructed using an Arima High Coverage Kit (Arima Genomics, CA, USA) according to the manufacturer’s instructions for mammalian cell lines (A160161 v01) and the Arima Library Prep Module (A160186 v01). Briefly, synchronised *B. dendrobatidis* cultures were filtered with a 10 µm cell strainer (pluriselect) to isolate zoospores, followed by *in situ* cell crosslinking, cell lysis chromatin digestion with 4 restriction enzymes (^GATC, G^ANTC, C^TNAG, T^TAA), biotin labelling, proximal chromatin DNA ligation and DNA purification. The resulting DNA was fragmented by sonication (Covaris E220) and enriched in biotinylated fragments followed by library preparation. The Hi-C library was subjected to paired-end sequencing with 150bp read lengths on an Illumina NovaSeq platform, resulting in 166 million read pairs (49.8 Gb, 2011X coverage).

### Genome assembly

The LongHam pipeline (long-read hybrid assembly merger) (Marcet-Houben et al 2022) was used to assemble ONT reads for *B. dendrobatidis* JEL423 (this study) and paired Illumina reads for *B. dendrobatidis* JEL423 (Farrer et al. 2013) to generate candidate primary assemblies. The hybrid *de novo* genome assembly that demonstrated the best mix of contiguity, completeness and number of telomeres was generated by the Maryland Super-Read Celera Assembler v.4.1.0 (MaSuRCA) (Zimin et al. 2013) with parameters “GRAPH_KMER_SIZE = auto, USE_LINKING_MATES = 0, LIMIT_JUMP_COVERAGE = 300, CA_PARAMETERS = cgwErrorRate=0.15, KMER_COUNT_THRESHOLD = 1, NUM_THREADS = 16, JF_SIZE = 480000000, SOAP_ASSEMBLY=0”. For polishing, Pilon v1.24 (Walker et al. 2014) was used with Illumina paired-end reads aligned to the draft assembly and indexed using BWA v.0.7.17 (Li 2013 Mar 16). The draft assembly and the corresponding alignments were then passed to Pilon v.1.24 (Walker et al. 2014) to call the consensus sequence.

Hi-C reads were aligned to the de novo assembly following the Arima mapping pipeline (https://github.com/ArimaGenomics/mapping_pipeline). Paired-end Illumina reads were first mapped independently to the genome sequence using BWA mem (Li 2013 Mar 16). Next, chimeric reads were filtered to only retain the 5’ side. Reads were then sorted, mapping quality filtered and paired, followed by removal of PCR duplicates. The final output is a single BAM file that contains the paired, 5’-filtered and duplicate-removed Hi-C reads, mapped to the reference sequence. Contigs were scaffolded with YaHS version 1.1 (Zhou et al. 2023). Mis-assemblies of contigs were manually corrected based using the resulting Hi-C contact map with Juicer Tools version 1.19.2 (Dudchenko et al. 2018) and Juicebox version 2.15 (Dudchenko et al. 2018). The resulting assembly has 18 scaffolds and 11 gaps. TGS-GapCloser version 1.1.1 (Xu et al. 2020) was used to fill gaps in the scaffolded assembly (using the long reads corrected with the short reads by Pilon v1.24 (Walker et al. 2014)). A total of 7 gaps were closed resulting in the final assembly of 18 scaffolds and 4 gaps. Scaffold ends were screened for telomeric sequences (TTAGGGn) with find_telomeric_repeats.py that is part of the LongHam pipeline (Marcet-Houben et al. 2022).

To calculate mapping rate, we aligned Illumina reads to the genome assembly using BWA v0.7.17 (Li 2013 Mar 16) and ONT long reads using minimap2 v2.28 (Li 2018). The rate was then calculated with SAMtools v1.20 flagstat (Li et al. 2009). To assess assembly completeness, we searched the *B. dendrobatidis* JEL423 genome sequence for a panel of core genes using Benchmarking Universal Single-Copy Orthologs (BUSCO) version 5.7.0 (Manni et al. 2021) specific to the fungi odb10 database.

### Genome annotation

The genome assembly was screened for repetitive sequence and masked using RepeatModeler2 (Flynn et al. 2020) and RepeatMasker version 4.1.5 (Smit et al.). Braker3 (Gabriel et al. 2024 Feb 29) was used to annotate the genome with esmode (for gene prediction with genome sequence data only) and softmasking enabled which combines GeneMark-ET version 3.68 (Lukashin and Borodovsky 1998) and AUGUSTUS version 3.5.0 (Stanke et al. 2006). EDTA version 2.2.0 (Ou et al. 2019) was used to annotate and classify transposable elements across the genome. We predicted the presence of signal peptides using SignalP4 (Petersen et al. 2011).

The protease composition of each chytrid was determined using top high scoring pairs from BLASTp searches (e-value < 1e-5) made to the file ‘pepunit.lib’, which is a non-redundant library of protein sequences of all the peptidases and peptidase inhibitors that are included in the MEROPS database (release 12.4) (Rawlings et al. 2016). All proteases with matches to M36 metalloproteases were aligned using MUSCLE v5.1.osx64 (Edgar 2022).

We constructed the gene trees with FastTree version 2.1.11 SSE3 (Price et al. 2009) with default parameters and visualised with FigTree v1.4.4 (http://tree.bio.ed.ac.uk/software/figtree/).

Crinklers and CBM18s were predicted based on BLASTp searches (e-value < 1e-5) to previously identified sequences (Farrer et al. 2017b). Two additional putative CBM18s (g4693 and g4694) were identified by HMMER v.3.1b2 (Eddy 2011) (hmmbuild the PFAM PF00187 and hmmscan all proteins). Interpro domain searches were used to identify and name domains. Completion of M36 metalloprotease and crinkler gene annotation was checked using tblastn v.2.15.0+ against our new *B. dendrobatidis* JEL423 genome assembly, and top HSPs and any overlapping predicted features were manually checked. Putative crinkler protein sequences were aligned using MUSCLE v5.1.osx64 (Edgar 2022) and gene trees constructed with FastTree version 2.1.11 SSE3 (Price et al. 2009).

### Orthology prediction, synteny, aneuploidy and phylogenetics

Orthologs were predicted between our new *B. dendrobatidis* JEL423 annotation and the previous (V2) assembly and annotation using the Synima pipeline (Farrer 2017) with OrthoMCL (Li et al. 2003). We also performed comparative genomics between *B. dendrobatidis* and six of its closest relatives (Wacker et al. 2023) that were downloaded from the MycoCosm portal of the US Department of Energy (DOE) Joint Genome Institute (JGI) (Grigoriev et al. 2014) (specifically, *Gorgonomyces haynaldii* MP57 (Amses et al. 2022), *Globomyces pollinis-pini* Arg68 (Amses et al. 2022), *Homolaphlyctis polyrhiza* JEL142 (Farrer 2016), *Batrachochytrium salamandrivorans* AMFP13 (GCA_002006685.2) (Wacker et al. 2023) and *Entophlyctis helioformis* JEL805 (Amses et al. 2022). Additionally, we included the genome for recently discovered *Polyrhizophydium stewartii* JEL0888 (GCA_027604665.1) (Simmons et al. 2021). The Synima pipeline outputs Orthogroups, which we divided into categories of interest, including single copy orthologs, which we used to construct phylogenetic trees. We generated a phylogenetic tree using 1,186 single copy orthologs. Single copy orthologs were aligned individually using MUSCLE version 5.1.osx64 (Edgar 2004) with default settings. We concatenated all alignments into a single FASTA and converted that into nexus format for phylogenetic analysis. We used the PROTCATWAG model for protein evolution with RAxML v.8.2.12 (Stamatakis 2014) with 1000 bootstraps.

Plotting ploidy and allele frequencies were performed using bespoke Perl scripts processing the data in pileup format (https://github.com/rhysf/allele_frequencies).

## Data availability

All ONT reads were uploaded to NCBI Sequence Read Achieve under project accession PRJNA1111108. All Hi-C reads were uploaded to NCBI Sequence Read Achieve under project accession PRJNA1111108. The annotated genome assembly for *B. dendrobatidis* JEL423 has been deposited to NCBI Genbank under project accession PRJNA1109278.

## Acknowledgements

We acknowledge funding from the MRC Centre for Medical Mycology at the University of Exeter (MR/N006364/2 and MR/V033417/1), and the NIHR Exeter Biomedical Research Centre. Additional work may have been undertaken by the University of Exeter Biological Services Unit. The views expressed are those of the author(s) and not necessarily those of the NIHR or the Department of Health. and Social Care. RAF is supported by a Wellcome Trust Career Development Award (225303/Z/22/Z). JH is supported with MRC funding (P76970). JG is supported by the MRC Doctoral Training Grant MR/W502649/1. EB is supported by a Sir Henry Dale Fellowship jointly funded by the Wellcome Trust and the Royal Society (211241/Z/18/Z); and the Lister Institute. We would like to thank Matthew Fisher, Theresa Wacker, Diana Tamayo, Qinxi Ma and Rahul Anand for useful discussions. We would like to thank Paul O’Neill and the Exeter Sequencing Service for their assistance with sequencing and base calling. This project utilised equipment funded by the UK Medical Research Council (MRC) Clinical Research Infrastructure Initiative (award number MR/M008924/1)

---

**Figure S1.** Counts of all predicted proteases based on a BLASTp search of the MEROPS database, comparing the number found in the previous assembly (V2) with the new assembly (V3).

**Figure S2.** M36 metalloproteases were predicted in the *B. dendrobatidis* JEL423 (V3) gene annotation based on BLASTp search of the MEROPS database. Tblastn against the genome identified additional high scoring pairs over genes highlighted in red. All genes identified by MEROPS BLASTP or Tblastn were aligned using MUSCLE and a tree constructed using FastTree (branch lengths indicate the mean number of nucleotide substitutions per site).

**Figure S3.** Predicted crinkling and necrosis (CRN) genes in B. dendrobatidis JEL423 and three chytrid relatives based on a BLASTp search to the *P. infestans* T30-4 as previously done (Farrer et al. 2017b).

**Table S1.** The number of telomere repeats identified on the 3; and 5’ ends of each scaffold of the new assembly.

## References

1. Abramyan J, Stajich JE. 2012. Species-specific chitin-binding module 18 expansion in the amphibian pathogen *Batrachochytrium dendrobatidis*. mBio. 3(3):e00150–00112. doi:10.1128/mBio.00150-12.

2. Amaro TMMM, Thilliez GJA, Motion GB, Huitema E. 2017. A Perspective on CRN Proteins in the Genomics Age: Evolution, Classification, Delivery and Function Revisited. Front Plant Sci. 8. doi:10.3389/fpls.2017.00099. [accessed 2024 Oct 7]. https://www.frontiersin.org/journals/plant-science/articles/10.3389/fpls.2017.00099/full.

3. Amses KR, Simmons DR, Longcore JE, Mondo SJ, Seto K, Jerônimo GH, Bonds AE, Quandt CA, Davis WJ, Chang Y, et al. 2022. Diploid-dominant life cycles characterize the early evolution of Fungi. Proc Natl Acad Sci. 119(36):e2116841119. doi:10.1073/pnas.2116841119.

4. Badet T, Tralamazza SM, Feurtey A, Croll D. 2024. Recent reactivation of a pathogenicity-associated transposable element is associated with major chromosomal rearrangements in a fungal wheat pathogen. Nucleic Acids Res. 52(3):1226–1242. doi:10.1093/nar/gkad1214.

5. Dudchenko O, Shamim MS, Batra SS, Durand NC, Musial NT, Mostofa R, Pham M, Hilaire BGS, Yao W, Stamenova E, et al. 2018. The Juicebox Assembly Tools module facilitates de novo assembly of mammalian genomes with chromosome-length scaffolds for under $1000. :254797. doi:10.1101/254797. [accessed 2024 Sep 16]. https://www.biorxiv.org/content/10.1101/254797v1.

6. Dupeyron M, Singh KS, Bass C, Hayward A. 2019. Evolution of Mutator transposable elements across eukaryotic diversity. Mob DNA. 10(1):12. doi:10.1186/s13100-019-0153-8.

7. Eddy SR. 2011. Accelerated profile HMM searches. PLoS Comput Biol. 7(10):e1002195. doi:10.1371/journal.pcbi.1002195.

8. Edgar RC. 2004. MUSCLE: a multiple sequence alignment method with reduced time and space complexity. BMC Bioinformatics. 5:113. doi:10.1186/1471-2105-5-113.

9. Edgar RC. 2022. Muscle5: High-accuracy alignment ensembles enable unbiased assessments of sequence homology and phylogeny. Nat Commun. 13(1):6968. doi:10.1038/s41467-022-34630-w.

10. Farrer R. 2016. Homolaphlyctis polyrhiza annotation GFF3. figshare. Dataset. 10.6084/m9.figshare.4291274.v2.

11. Farrer RA. 2017. Synima: a synteny imaging tool for annotated genome assemblies. BMC Bioinformatics. 18(1):507. doi:10.1186/s12859-017-1939-7.

12. Farrer RA. 2019. Batrachochytrium salamandrivorans. Trends Microbiol. 27(10):892–893. doi:10.1016/j.tim.2019.04.009.

13. Farrer RA. 2021. HaplotypeTools: a toolkit for accurately identifying recombination and recombinant genotypes. BMC Bioinformatics. 22(1):560. doi:10.1186/s12859-021-04473-1.

14. Farrer RA, Henk DA, Garner TWJ, Balloux F, Woodhams DC, Fisher MC. 2013. Chromosomal copy number variation, selection and uneven rates of recombination reveal cryptic genome diversity linked to pathogenicity. PLOS Genet. 9(8):e1003703. doi:10.1371/journal.pgen.1003703.

15. Farrer RA, Martel A, Verbrugghe E, Abouelleil A, Ducatelle R, Longcore JE, James TY, Pasmans F, Fisher MC, Cuomo CA. 2017a. Genomic innovations linked to infection strategies across emerging pathogenic chytrid fungi. Nat Commun. 8:14742. doi:10.1038/ncomms14742.

16. Farrer RA, Martel A, Verbrugghe E, Abouelleil A, Ducatelle R, Longcore JE, James TY, Pasmans F, Fisher MC, Cuomo CA. 2017b. Genomic innovations linked to infection strategies across emerging pathogenic chytrid fungi. Nat Commun. 8(1):14742. doi:10.1038/ncomms14742.

17. Farrer RA, Weinert LA, Bielby J, Garner TWJ, Balloux F, Clare F, Bosch J, Cunningham AA, Weldon C, du Preez LH, et al. 2011a. Multiple emergences of genetically diverse amphibian- infecting chytrids include a globalized hypervirulent recombinant lineage. Proc Natl Acad Sci U S A. 108(46):18732–18736. doi:10.1073/pnas.1111915108.

18. Farrer RA, Weinert LA, Bielby J, Garner TWJ, Balloux F, Clare F, Bosch J, Cunningham AA, Weldon C, du Preez LH, et al. 2011b. Multiple emergences of genetically diverse amphibian- infecting chytrids include a globalized hypervirulent recombinant lineage. Proc Natl Acad Sci. 108(46):18732–18736. doi:10.1073/pnas.1111915108.

19. Fisher MC, Garner TWJ. 2020. Chytrid fungi and global amphibian declines. Nat Rev Microbiol. 18(6):332–343. doi:10.1038/s41579-020-0335-x.

20. Flynn JM, Hubley R, Goubert C, Rosen J, Clark AG, Feschotte C, Smit AF. 2020. RepeatModeler2 for automated genomic discovery of transposable element families. Proc Natl Acad Sci U S A. 117(17):9451–9457. doi:10.1073/pnas.1921046117.

21. Gabriel L, Brůna T, Hoff KJ, Ebel M, Lomsadze A, Borodovsky M, Stanke M. 2024 Feb 29. BRAKER3: Fully automated genome annotation using RNA-seq and protein evidence with GeneMark-ETP, AUGUSTUS and TSEBRA. BioRxiv Prepr Serv Biol.:2023.06.10.544449. doi:10.1101/2023.06.10.544449.

22. Gao X, Hou Y, Ebina H, Levin HL, Voytas DF. 2008. Chromodomains direct integration of retrotransposons to heterochromatin. Genome Res. 18(3):359–369. doi:10.1101/gr.7146408.

23. Greenspan SE, Lambertini C, Carvalho T, James TY, Toledo LF, Haddad CFB, Becker CG. 2018. Hybrids of amphibian chytrid show high virulence in native hosts. Sci Rep. 8(1):9600. doi:10.1038/s41598-018-27828-w.

24. Grigoriev IV, Nikitin R, Haridas S, Kuo A, Ohm R, Otillar R, Riley R, Salamov A, Zhao X, Korzeniewski F, et al. 2014. MycoCosm portal: gearing up for 1000 fungal genomes. Nucleic Acids Res. 42(Database issue):D699-704. doi:10.1093/nar/gkt1183.

25. Hoencamp C, Dudchenko O, Elbatsh AMO, Brahmachari S, Raaijmakers JA, van Schaik T, Sedeño Cacciatore Á, Contessoto VG, van Heesbeen RGHP, van den Broek B, et al. 2021. 3D genomics across the tree of life reveals condensin II as a determinant of architecture type. Science. 372(6545):984–989. doi:10.1126/science.abe2218.

26. Joneson S, Stajich JE, Shiu S-H, Rosenblum EB. 2011. Genomic transition to pathogenicity in chytrid fungi. PLOS Pathog. 7(11):e1002338. doi:10.1371/journal.ppat.1002338.

27. Kalinka E, Brody SM, Swafford AJM, Medina EM, Fritz-Laylin LK. 2024. Genetic transformation of the frog-killing chytrid fungus *Batrachochytrium dendrobatidis*. Proc Natl Acad Sci U S A. 121(4):e2317928121. doi:10.1073/pnas.2317928121.

28. Kapitonov VV, Jurka J. 2006. Self-synthesizing DNA transposons in eukaryotes. Proc Natl Acad Sci. 103(12):4540–4545. doi:10.1073/pnas.0600833103.

29. Li H. 2013 Mar 16. Aligning sequence reads, clone sequences and assembly contigs with BWA-MEM. ArXiv13033997 Q-Bio. [accessed 2018 May 29]. http://arxiv.org/abs/1303.3997.

30. Li H. 2018. Minimap2: pairwise alignment for nucleotide sequences. Bioinformatics. 34(18):3094–3100. doi:10.1093/bioinformatics/bty191.

31. Li H, Handsaker B, Wysoker A, Fennell T, Ruan J, Homer N, Marth G, Abecasis G, Durbin R, 1000 Genome Project Data Processing Subgroup. 2009. The Sequence Alignment/Map format and SAMtools. Bioinforma Oxf Engl. 25(16):2078–2079. doi:10.1093/bioinformatics/btp352.

32. Li L, Stoeckert CJ, Roos DS. 2003. OrthoMCL: identification of ortholog groups for eukaryotic genomes. Genome Res. 13(9):2178–2189. doi:10.1101/gr.1224503.

33. Liu P, Stajich JE. 2015. Characterization of the Carbohydrate Binding Module 18 gene family in the amphibian pathogen *Batrachochytrium dendrobatidis*. Fungal Genet Biol FG B. 77:31–39. doi:10.1016/j.fgb.2015.03.003.

34. Longcore JE, Pessier AP, Nichols DK. 1999. *Batrachochytrium Dendrobatidis* gen. et sp. nov., a Chytrid Pathogenic to Amphibians. Mycologia. 91(2):219–227. doi:10.2307/3761366.

35. Lukashin AV, Borodovsky M. 1998. GeneMark.hmm: new solutions for gene finding. Nucleic Acids Res. 26(4):1107–1115.

36. Manni M, Berkeley MR, Seppey M, Zdobnov EM. 2021. BUSCO: Assessing Genomic Data Quality and Beyond. Curr Protoc. 1(12):e323. doi:10.1002/cpz1.323.

37. Marcet-Houben M, Alvarado M, Ksiezopolska E, Saus E, de Groot PWJ, Gabaldón T. 2022. Chromosome-level assemblies from diverse clades reveal limited structural and gene content variation in the genome of *Candida glabrata*. BMC Biol. 20:226. doi:10.1186/s12915-022-01412-1.

38. Muszewska A, Hoffman-Sommer M, Grynberg M. 2011. LTR Retrotransposons in Fungi. PLoS ONE. 6(12):e29425. doi:10.1371/journal.pone.0029425.

39. O’Hanlon SJ, Rieux A, Farrer RA, Rosa GM, Waldman B, Bataille A, Kosch TA, Murray KA, Brankovics B, Fumagalli M, et al. 2018. Recent Asian origin of chytrid fungi causing global amphibian declines. Science. 360(6389):621–627. doi:10.1126/science.aar1965.

40. Ou S, Su W, Liao Y, Chougule K, Agda JRA, Hellinga AJ, Lugo CSB, Elliott TA, Ware D, Peterson T, et al. 2019. Benchmarking transposable element annotation methods for creation of a streamlined, comprehensive pipeline. Genome Biol. 20(1):275. doi:10.1186/s13059-019-1905-y.

41. Petersen TN, Brunak S, von Heijne G, Nielsen H. 2011. SignalP 4.0: discriminating signal peptides from transmembrane regions. Nat Methods. 8(10):785–786. doi:10.1038/nmeth.1701.

42. Price MN, Dehal PS, Arkin AP. 2009. FastTree: computing large minimum evolution trees with profiles instead of a distance matrix. Mol Biol Evol. 26(7):1641–1650. doi:10.1093/molbev/msp077.

43. Rawlings ND, Barrett AJ, Finn R. 2016. Twenty years of the MEROPS database of proteolytic enzymes, their substrates and inhibitors. Nucleic Acids Res. 44(D1):D343–350. doi:10.1093/nar/gkv1118.

44. Rosenblum EB, James TY, Zamudio KR, Poorten TJ, Ilut D, Rodriguez D, Eastman JM, Richards-Hrdlicka K, Joneson S, Jenkinson TS, et al. 2013. Complex history of the amphibian- killing chytrid fungus revealed with genome resequencing data. Proc Natl Acad Sci U S A. 110(23):9385–9390. doi:10.1073/pnas.1300130110.

45. Scheele BC, Pasmans F, Skerratt LF, Berger L, Martel A, Beukema W, Acevedo AA, Burrowes PA, Carvalho T, Catenazzi A, et al. 2019. Amphibian fungal panzootic causes catastrophic and ongoing loss of biodiversity. Science. 363(6434):1459–1463. doi:10.1126/science.aav0379.

46. Schwessinger B. 2019 May 17. High quality DNA from Fungi for long read sequencing e.g. PacBio. protocols.io. doi:10.17504/protocols.io.2yfgftn. [accessed 2020 Aug 20]. https://www.protocols.io/view/high-quality-dna-from-fungi-for-long-read-sequenci-2yfgftn.

47. Simmons DR, Longcore JE, James TY. 2021. *Polyrhizophydium stewartii*, the first known rhizomycelial genus and species in the Rhizophydiales, is closely related to Batrachochytrium. Mycologia. 113(3):684–690. doi:10.1080/00275514.2021.1885206.

48. Smit A, Hubley R, Green P. RepeatMasker Open-4.0. [accessed 2017 Mar 24]. http://www.repeatmasker.org/.

49. Stamatakis A. 2014. RAxML version 8: a tool for phylogenetic analysis and post-analysis of large phylogenies. Bioinformatics. 30(9):1312–1313. doi:10.1093/bioinformatics/btu033.

50. Stanke M, Keller O, Gunduz I, Hayes A, Waack S, Morgenstern B. 2006. AUGUSTUS: *ab initio* prediction of alternative transcripts. Nucleic Acids Res. 34(Web Server issue):W435–W439. doi:10.1093/nar/gkl200.

51. The IUCN Red List of Threatened Species. 2022. IUCN Red List Threat Species. [accessed 2024 Mar 5]. https://www.iucnredlist.org/en.

52. Wacker T, Helmstetter N, Studholme DJ, Farrer RA. 2024. Genome variation in the *Batrachochytrium* pathogens of amphibians. PLOS Pathog. 20(5):e1012218. doi:10.1371/journal.ppat.1012218.

53. Wacker T, Helmstetter N, Wilson D, Fisher MC, Studholme DJ, Farrer RA. 2023. Two-speed genome evolution drives pathogenicity in fungal pathogens of animals. Proc Natl Acad Sci U S A. 120(2):e2212633120. doi:10.1073/pnas.2212633120.

54. Walker BJ, Abeel T, Shea T, Priest M, Abouelliel A, Sakthikumar S, Cuomo CA, Zeng Q, Wortman J, Young SK, et al. 2014. Pilon: an integrated tool for comprehensive microbial variant detection and genome assembly improvement. PloS One. 9(11):e112963. doi:10.1371/journal.pone.0112963.

55. Webb RJ, Vu AL, Skerratt LF, Berger L, Andino FDJ, Robert J. 2024. Stable *in vitro* fluorescence for enhanced live imaging of infection models for *Batrachochytrium dendrobatidis*. PLOS ONE. 19(8):e0309192. doi:10.1371/journal.pone.0309192.

56. Xu M, Guo L, Gu S, Wang O, Zhang R, Peters BA, Fan G, Liu X, Xu X, Deng L, et al. 2020. TGS- GapCloser: A fast and accurate gap closer for large genomes with low coverage of error- prone long reads. GigaScience. 9(9):giaa094. doi:10.1093/gigascience/giaa094.

57. Yacoub MN, Stajich JE. 2024. Comparative genomics reveals intra and inter species variation in the pathogenic fungus Batrachochytrium dendrobatidis. :2024.01.24.576925. doi:10.1101/2024.01.24.576925. [accessed 2024 Jan 25]. https://www.biorxiv.org/content/10.1101/2024.01.24.576925v1.

58. Zhou C, McCarthy SA, Durbin R. 2023. YaHS: yet another Hi-C scaffolding tool. Bioinformatics. 39(1):btac808. doi:10.1093/bioinformatics/btac808.

59. Zimin AV, Marçais G, Puiu D, Roberts M, Salzberg SL, Yorke JA. 2013. The MaSuRCA genome assembler. Bioinforma Oxf Engl. 29(21):2669–2677. doi:10.1093/bioinformatics/btt476.

